# Detecting quantitative trait loci and exploring chromosomal pairing in autopolyploids using *polyqtlR*

**DOI:** 10.1101/2021.06.28.450123

**Authors:** Peter M. Bourke, Roeland E. Voorrips, Christine A. Hackett, Geert van Geest, Johan H. Willemsen, Paul Arens, Marinus J. M. Smulders, Richard G. F. Visser, Chris Maliepaard

## Abstract

**Motivation:** The investigation of quantitative trait loci (QTL) is an essential component in our understanding of how organisms vary phenotypically. However, many important crop species are polyploid (carrying more than two copies of each chromosome), requiring specialised tools for such analyses. Moreover, deciphering meiotic processes at higher ploidy levels is not straightforward, but is necessary to understand the reproductive dynamics of these species, or uncover potential barriers to their genetic improvement.

**Results:** Here we present *polyqtlR*, a novel software tool to facilitate such analyses in (auto)polyploid crops. It performs QTL interval mapping in F_1_ populations of outcrossing polyploids of any ploidy level using identity-by-descent (IBD) probabilities. The allelic composition of discovered QTL can be explored, enabling favourable alleles to be identified and tracked in the population. Visualisation tools within the package facilitate this process, and options to include genetic co-factors and experimental factors are included. Detailed information on polyploid meiosis including prediction of multivalent pairing structures, detection of preferential chromosomal pairing and location of double reduction events can be performed.

**Availability and implementation:** *polyqtlR* is freely available from the Comprehensive R Archive Network (CRAN) at http://cran.r-project.org/package=polyqtlR.

**Supplementary information:** Supplementary data are available

## Introduction

Polyploids, which carry more than two copies of each chromosome, are an important group of organisms that occur widely among plant species, including several domesticated crops (Salman-Minkov et al., 2016). Many theories to explain their prevalence among crop species have been proposed, identifying features which may have appealed to early farmers in their domestication of wild species. Such features include their larger organs such as tubers, fruits or flowers (the so-called “gigas” effect) (Sattler et al., 2016), phenotypic novelty (Udall and Wendel, 2006), their ability to be clonally propagated (Herben et al., 2017), increased seedling and juvenile vigour (Levin, 1983) and the possibility of seedlessness which accompanies odd-numbered ploidies (Bradshaw, 2016). From a functional perspective, these features may be associated with factors such as increased heterosis (Comai, 2005), a greater level of genomic plasticity (te Beest et al., 2011) or a masking effect of deleterious alleles (Renny-Byfield and Wendel, 2014). It is currently believed that all flowering plants have experienced at least one whole genome duplication (WGD) during the course of their evolution, with many lineages undergoing multiple rounds of WGD followed by re-diploidisation (Vanneste et al., 2014). Polyploidy may also be induced deliberately (through species hybridisation with associated unreduced gametes, or through the use of some chemical cell division inhibitor such as colchicine (Blakeslee and Avery, 1937)), often to combine properties of parents that could not otherwise be crossed (Van Tuyl and Lim, 2003), or to benefit from some of the other advantages listed above.

Polyploids are generally divided into two groups: autopolyploids, with multiple copies of the same homologous chromosomes derived from a single progenitor species, and allopolyploids, with multiple copies of homoeologous chromosomes from multiple progenitor species that continue to pair and recombine within but not between homoeologues. Allopolyploids are said to exhibit “disomic” inheritance *i*.*e*. genetically-speaking they are equivalent to diploids.

Autopolyploids are, on the other hand, genetically distinct from allopolyploids in that they exhibit “polysomic” inheritance, a property that emerges from random pairing and recombination between homologues during meiosis. As most software and methodology for genetic analyses has traditionally been developed for diploid organisms, progress in autopolyploid breeding and research has been slower. In recent years this has gradually changed, as more tools become available for autopolyploids too (Bourke et al., 2018c).

One of the greatest difficulties in autopolyploid cultivation and breeding is the constant reshuffling of alleles in each generation, a consequence of polysomic inheritance. Breeders would like to be able to identify genomic regions that contribute favourable alleles to a particular trait of interest (quantitative trait loci (QTL)) and predict which offspring in a population carry favourable combinations of parental alleles. Understanding how genes and their alleles are transmitted from one generation to the next, or identifying potential barriers to recombination that might restrict allelic combinations from arising can provide insights into designing crosses and identifying favourable offspring from these crosses. The use of genomic information can greatly assist in these efforts. Particularly for polyploid species, specialised software tools are required for this purpose.

Polyploid genotyping involves the estimation of dosage (counts of the alternative allele at a polymorphic site, usually bi-allelic single nucleotide polymorphisms (SNPs)). In an autotetraploid for example, the possible dosages range from nulliplex (0 copies of the alternative allele), simplex (1 copy), duplex (2 copies), triplex (3 copies) to quadruplex (4 copies). The assignment of marker dosage in polyploids is a non-trivial problem in itself, but there are an increasing number of possibilities for achieving this using dedicated software (Voorrips et al., 2011;Serang et al., 2012;Carley et al., 2017;Gerard et al., 2018;Pereira et al., 2018;Clark et al., 2019;Zych et al., 2019).

Identity-by-descent (IBD) probabilities are the inheritance probabilities of parental alleles in a population of related genotypes (either bi-parental or multi-parental), and they can be exploited both for QTL mapping and to accurately interpret parental meiosis and inheritance patterns. Hidden Markov Models (HMM) have previously been applied to estimate these inheritance probabilities for polyploids (Hackett et al., 2013;Zheng et al., 2016;Mollinari and Garcia, 2019;Zheng et al., 2020), and have been shown to be robust against common issues such as genotyping errors or local ordering issues in the underlying linkage maps (Zheng et al., 2016). Of the currently available methods, both TetraOrigin and polyOrigin include a fully generalised polysomic model with the possibility of including multivalents in the model of parental meiosis (Zheng et al., 2016;Zheng et al., 2020). These packages are, however, currently aimed at tetraploid species. MAPpoly implements a HMM to estimate IBD probabilities that can be applied for all even ploidy levels, but assumes bivalent pairing only (Mollinari and Garcia, 2019). However, autopolyploids carry homologous chromosomes that often pair during meiosis in more complex structures called multivalents, associations of more than two homologues (generally only even numbers are considered viable). In particular, the phenomenon of double reduction, a possible product of multivalent pairing where both copies of a segment of sister chromatids are passed on to an offspring (Bourke et al., 2015), is ignored. The overall impact of omitting double reduction from the model used for QTL analysis has previously been shown to be relatively minor in a QTL analysis that does not account for it (Bourke et al., 2019), but double reduction events at specific loci may have important breeding implications (*e*.*g*. an offspring carrying a double copy of a favourable allele at that locus). For higher ploidy levels (6x and higher), HMM approaches may lead to computational bottlenecks (Mollinari and Garcia, 2019). Alternative approaches to estimate IBD probabilities have been proposed (Bourke, 2014) and although less accurate, have the advantage of being computationally tractable at higher ploidy levels and have previously been successfully used in the analysis of several traits in hexaploid chrysanthemum for example (van Geest et al., 2017a).

Apart from their application in QTL mapping, identity-by-descent probabilities provide a powerful approach to reconstruct meiotic processes and identify recombination events in polyploid individuals. This latter point can be exploited to address the issue of genotyping errors. They also yield insights into potential preferential chromosomal pairing, which is increasingly being acknowledged as a feature of many polyploid species that were previously assumed to be either purely auto- or allopolyploid (Bourke et al., 2017;Leal-Bertioli et al., 2018). In this paper we describe the features of *polyqtlR*, a novel R package (R Core Team, 2020) for QTL mapping in both auto- and allopolyploid species which addresses many of the complexities of polyploid inheritance mentioned above. Estimation of identity-by-descent probabilities under a full polysomic model (including multivalents and double reduction) is performed for autotriploid, autotetraploid and autohexaploid F_1_ populations, while IBD probabilities of diploids and allopolyploids are estimated using a diploid HMM. Alternatively, a computationally-efficient but approximate method for IBD estimation suitable for all ploidy levels (allo- and auto-) is implemented in *polyqtlR*. With these IBD probabilities, a range of applications are available, for QTL discovery and exploration as well as investigation of meiotic processes and patterns of recombination across the genome.

## Materials and Methods

### Input data

*polyqtlR* requires as input dosage-scored marker information with an accompanying phased linkage map from an F_1_ population. Dosage scores can either be discrete or probabilistic (*i*.*e*. the probabilities of each of the dosage classes from 0 to *ploidy* for each individual at a marker), while phased linkage maps can be generated using software such as TetraploidSNPMap (Hackett et al., 2017), polymapR (Bourke et al., 2018b) or MAPpoly (Mollinari and Garcia, 2019). For hexaploid populations, only polymapR or MAPpoly are currently suitable, while polymapR is the only software that can also map odd-numbered ploidies such as triploid populations (Bourke et al., 2018b). In the case of tetraploids for which a marker order is already known, parental map phase and IBD probabilities can also be estimated using TetraOrigin or polyOrigin (Zheng et al., 2016;Zheng et al., 2020).

### Modelling autopolyploid meiosis

#### Hidden Markov Model

The methodology behind the estimation of offspring IBD probabilities was originally developed for tetraploid populations (Zheng et al., 2016) but we have extended the approach to a range of commonly-encountered ploidy levels (2x, 3x, 4x and 6x). Details are contained in Supplemental Methods SM1.

#### Heuristic model

An algorithm for approximating IBD probabilities without using Hidden Markov Models is also implemented in *polyqtlR*. This uses an approach originally described in Bourke et al. (2014) and re-implemented by van Geest et al. (2017). Details are contained in Supplementary Methods SM2.

Finally, IBD probabilities may be interpolated at a regular grid of positions using cubic splines (by default at 1 cM spacings).

#### Form of the QTL model

The IBD-based QTL analysis uses a linear regression on the parental homologue probabilities, broadly similar to the weighted regression model proposed by Kempthorne (Kempthorne, 1957) and implemented in the TetraploidSNPMap software (Hackett et al., 2013;Hackett et al., 2014;Hackett et al., 2017). For a tetraploid, the form of the model is:

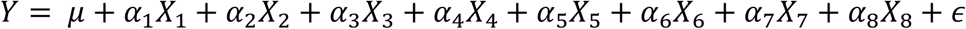

Here, the *X*_*i*_ are inheritance probabilities for each parental homologue (1-4 for parent 1, 5-8 for parent 2 in a tetraploid), which range from 0 ≤ *X*_*i*_ ≤ 1 when bivalent-only pairing is assumed. In the case where multivalents are also permitted in the meiotic model, more than one copy of a parental homologue can be inherited through the process of double reduction, in which case *X*_*i*_ are the total sum of inheritance probabilities for each parental homologue, with 0 ≤ *X*_*i*_ ≤ 2. In the context of a tetraploid, it can be generally assumed *X*_1_ + *X*_2_ + *X*_3_ + *X*_4_ = 2 and *X*_5_ + *X*_6_ + *X*_7_ + *X*_8_ = 2 (*i*.*e*. both parents contribute an equal number of chromosomes ot an offspring). Eliminating *X*_1_ and *X*_5_ by substituting these expressions into the previous equation (to remove collinearity) leads to the following model:

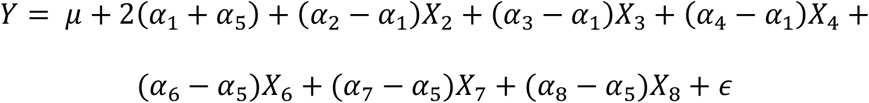

This can be re-written as:

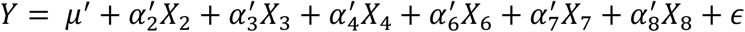

where *μ*′ is the adjusted intercept (*μ*′ = *μ* + 2(*α* _1_ + *α* _5_)), 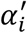 are the adjusted regression coefficients (*e*.*g*. 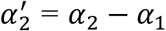) and *ϵ* is the residual term. For a hexaploid, the model includes ten of the twelve parental homologues (van Geest et al., 2017a) *etc*.

A single marker analysis option is also included in the package, in which a genome-wide scan is performed by fitting the following additive model at each marker position:

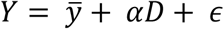

where *Y* is the vector of phenotypes, *α* is the vector of marker scores, 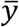 is the overall mean and *ϵ* the residuals.

If experimental factors (loosely termed “blocks” here, although they could correspond to different years, environments *etc*.) are included, they are first fitted (*Y* ∼ *Blocks*) after which the residuals are used to perform the genome-wide QTL scan. Missing phenotypes are imputed using fitted block effects and non-missing phenotype scores for that individual in other blocks. By default, at least 50% observations are required for imputation (*e*.*g*. minimum 2 out of 3 phenotypes non-missing for that individual in a 3-block situation). Estimating BLUEs (using a linear mixed model with genotypes as fixed effects (Pinheiro et al., 2017)) for block-corrected trait values can speed up the analyses, particularly when estimating significance thresholds.

The sum of squared residuals (*RSS*_*1*_) is recorded from the ANOVA table (for both IBD-based and single marker approaches) and used to calculate the logarithm of odds ratio (LOD) score as follows (Broman and Sen, 2009):

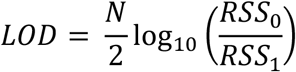

where *N* is the population size, and *RSS*_*0*_ is the residual sum of squares under the Null (no QTL) Model. In cases where large-effect QTL are present and segregating in a population, it can be advantageous to reduce the level of background noise at other loci by accounting first for the major QTL and running an analysis on the QTL-corrected phenotypes. Such an approach has previously been termed multiple QTL mapping (Jansen, 1992;Jansen, 1993). In *polyqtlR*, we follow a similar approach to correct for genetic co-factors, either by supplying the name of a marker closely linked to the major QTL peak, or the QTL peak position from the genome-wide scan (usually performed at regular intervals for efficiency). There is no limit to the number of co-factors that can be added, but a parsimonious analysis with only significant QTL as genetic co-factors is recommended (to avoid issues of collinearity). Automatic fitting of genetic co-factors is also implemented, fitting all possible combinations of initially detected QTL exceeding the significance threshold as co-factors (*i*.*e*. for QTL q_1_, q_2_, …. q_n_, all co-factor models 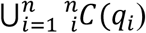 are tested, where 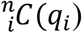 denotes all *i*-wise combinations of QTL for *i* ∈ [1,2, …, *n*]). Following this, a set of positions that individually maximised the threshold-adjusted LOD scores within the genetic region associated with each QTL locus are identified (a QTL locus is by default assumed to be no smaller than a 20 cM interval – *i*.*e*. this is the smallest assumed resolution between independent QTL that could occur). These new set of positions are then fed back into the same procedure to refine the estimates of QTL position and threshold-corrected significance, with positions that maximise the threshold-corrected LOD score being selected. . Internally, the QTL model described above for IBD probabilities is initially fitted at the supplied position(s) and the residuals are saved to replace the vector of phenotype values in the QTL scan. Note that when blocks or genetic co-factors are included, they form part of the Null Model in the calculation of LOD scores.

The percentage of phenotypic variance explained at a single position is estimated by

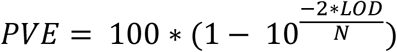

In the case of a multi-QTL model, the PVE is estimated using 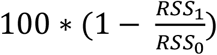, where *RSS*_*1*_ is now recorded from the fitted (multi-) QTL model and *RSS*_*0*_ from the no-QTL model (Broman and Sen, 2009).

Approximate significance thresholds are determined using Permutation Tests (Churchill and Doerge, 1994). The number of permutations *N*_*p*_ and the approximate Type I error rate α can be specified. By default *N*_*p*_ = 1000 permutations of trait values are performed, after which the maximum genome-wide LOD scores are recorded from each of the *N*_*p*_ genome-wide scans. The 100*(1 - α) percentile of the ordered LOD scores is taken as an approximate 100*(1 - α) % significance threshold (by default α = 0.05). Chromosome-specific thresholds can be generated by restricting the input to the chromosome(s) of interest, if so desired.

### Exploration of QTL configuration and mode of action

One of the advantages of an IBD-based analysis over single-marker methods is the ability to explore QTL peak positions to determine the most likely QTL configuration (the parental origin of QTL alleles that have an effect on the phenotype), their mode of action (additive / dominant) and the effect sizes (both positive and negative) of specific parental alleles. A range of QTL models can be compared in *polyqtlR* using the Bayesian Information Criterion (BIC) (Schwarz, 1978) as previously proposed (Hackett et al., 2014). Homologue-specific effects can be visualised around QTL peaks, aiding in the interpretation of the most likely predicted QTL configuration.

### Genotypic Information Coefficient (GIC)

The Genotypic Information Coefficient (GIC) is a convenient measure of the precision of our knowledge on the composition of parental alleles carried by each offspring individual at a particular position, averaged across the mapping population. This is visualised in a similar manner to QTL profile plots, providing an overview of the genome-wide information landscape in the population in the vicinity of detected (or indeed expected but undetected) QTL.

The GIC of homologue *j* (1 ≤ *j* ≤ *ploidy* + *ploidy*2) at each position is calculated from the IBD probabilities using the formula:

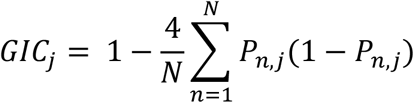

where *ploidy* and *ploidy*2 refer to the ploidy levels of the two parents, *N* is the population size and *P*_*n,j*_ is the probability that individual *n* inherited homologue *j* at that position (this is a generalisation of the GIC measure used in MapQTL (Van Ooijen, 1992;Van Ooijen, 2009)); for a derivation see Appendix I of Bourke *et al*. (2019). When multivalents are included in the HMM, only offspring predicted to have come from bivalent-only meioses for that linkage group are used in the calculation.

### Polyploid meiosis

Several aspects of polyploid meiosis can be investigated using IBD probabilities. These include detecting signatures of multivalent pairing structures in tetraploid and hexaploid parents, determining rates of double reduction, identifying recombinations from cross-overs, and looking for deviations from random pairing in meiosis, a feature associated with autopolyploidy (polysomic inheritance). Non-random chromosomal pairing, also called “preferential pairing” (Bourke et al., 2017) can be detected across a population using counts of bivalent pairing structures. Using the HMM method of IBD estimation, each valency (homologue pairing configuration) has an associated posterior probability. Deviations from random pairing are tested per parental chromosome using a chi-square test on the counts of predicted pairings.

### Cross-overs, errors and linkage map curation

Recombinations from cross-overs are detected given a predicted pairing by looking for regions in which the inheritance probabilities of pairing parental homologues switch from one homologue to the other. A threshold probability is defined (by default 0.4) to identify meiotic pairing patterns that were clearly predicted or “plausible” using the HMM and screen out those that were ambiguous. Within the context of bivalent pairing, each individual has an associated inheritance probability (IBD) for homologues in each bivalent pair. A recombination break-point is defined as a point at which the difference in inheritance probabilities of such pairing homologues changes sign. Its position is taken as the midpoint between the flanking positions for which such a switch-over in inheritance probabilities occurred. Individuals showing unexpectedly high numbers of recombinations can be identified and removed. One of the input parameters in the HMM method of IBD estimation is the error prior *ε*, the genome-wide error rate in the offspring genotypes. With high-quality data, error priors of the order 0.01 – 0.05 are reasonable, while for poorer-quality data a higher error prior may be required. If IBD probabilities are estimated using a suitably high error prior (*ε* = 0.2, say), spurious recombinations from genotyping errors are suppressed, in which case IBD parental homologue probabilities can be used to directly re-impute marker genotype dosages with the function impute_dosages. For each individual, the imputed dosage of individual *j* at marker *n* on a certain linkage group is given by:

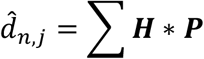

where ***H*** is the (*ploidy1* + *ploidy2*) x 1 vector of parental homologue probabilities of that individual at that marker position (*ploidy1* and *ploidy2* being the ploidy levels of parent 1 and parent 2, respectively), ***P*** is the (*ploidy1* + *ploidy2*) x 1 vector of parental phase coded in 0 and 1 notation (1 for presence, 0 for absence) and ***H*** * ***P*** is their element-wise product. This operation generally leads to non-integer dosage values, and so 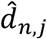 is rounded to the nearest integer. If the absolute value of the difference between the exact and rounded values exceeds a user-defined rounding error threshold (by default 0.05), the imputed dosage is set to missing.

## Results

We demonstrate the capabilities of *polyqtlR* with a number of example applications. We first analysed an example trait in both tetraploid and hexaploid material and compared our results to those generated using TetraploidSNPMap (Hackett et al., 2017) and QTLpoly (Pereira et al., 2020). We then used the package to dissect the meiotic patterns of a hexaploid chrysanthemum population (van Geest et al., 2017a). Finally, we performed some tests to quantify the accuracy and computational performance of the IBD estimation module in the package.

### 1. QTL detection

A tetraploid cut rose (*Rosa* x *hybrida*) dataset for the morphological trait ‘stem prickles’ using a previously published linkage map (Bourke et al., 2017) and phenotypic data collected in different growing environments (Bourke et al., 2018a) was analysed with *polyqtlR*, QTLpoly and TetraploidSNPMap. The first genome-wide scan for QTL using *polyqtlR* detected four putative QTL on LG 2, 3, 4 and 6 (Figure 1). By fitting various combinations of QTL as co-factors, we found that the significance of the LG 3, 4 and 6 peaks could be increased, while the peak on LG 2 dropped in significance upon the inclusion of co-factors. Different co-factor combinations were tested for each QTL. The analysis that resulted in the highest threshold-adjusted LOD score for that peak was used to estimate the QTL position. The four-QTL model was found to explain 54% of the phenotypic variation (PVE), while the best three- and two-QTL models explained 49% and 43% of the phenotypic variation, respectively. TetraploidSNPMap predicted a three-QTL model, detecting the same QTL on LG 3, 4 and 6, while a putative position on LG 2 failed to reach the genome-wide significance threshold. QTLpoly predicted a two-QTL model, detecting peaks on LG 3 and 4. A putative QTL was initially detected on LG 6 in the forward search (sig.fwd = 0.01), but was removed in the subsequent backward elimination step (sig.bwd = 0.0001). The major QTL on LG 3 and 4 have also been reported in previous studies in diploid rose populations (Crespel et al., 2002;Linde et al., 2006).

**Figure 1.**
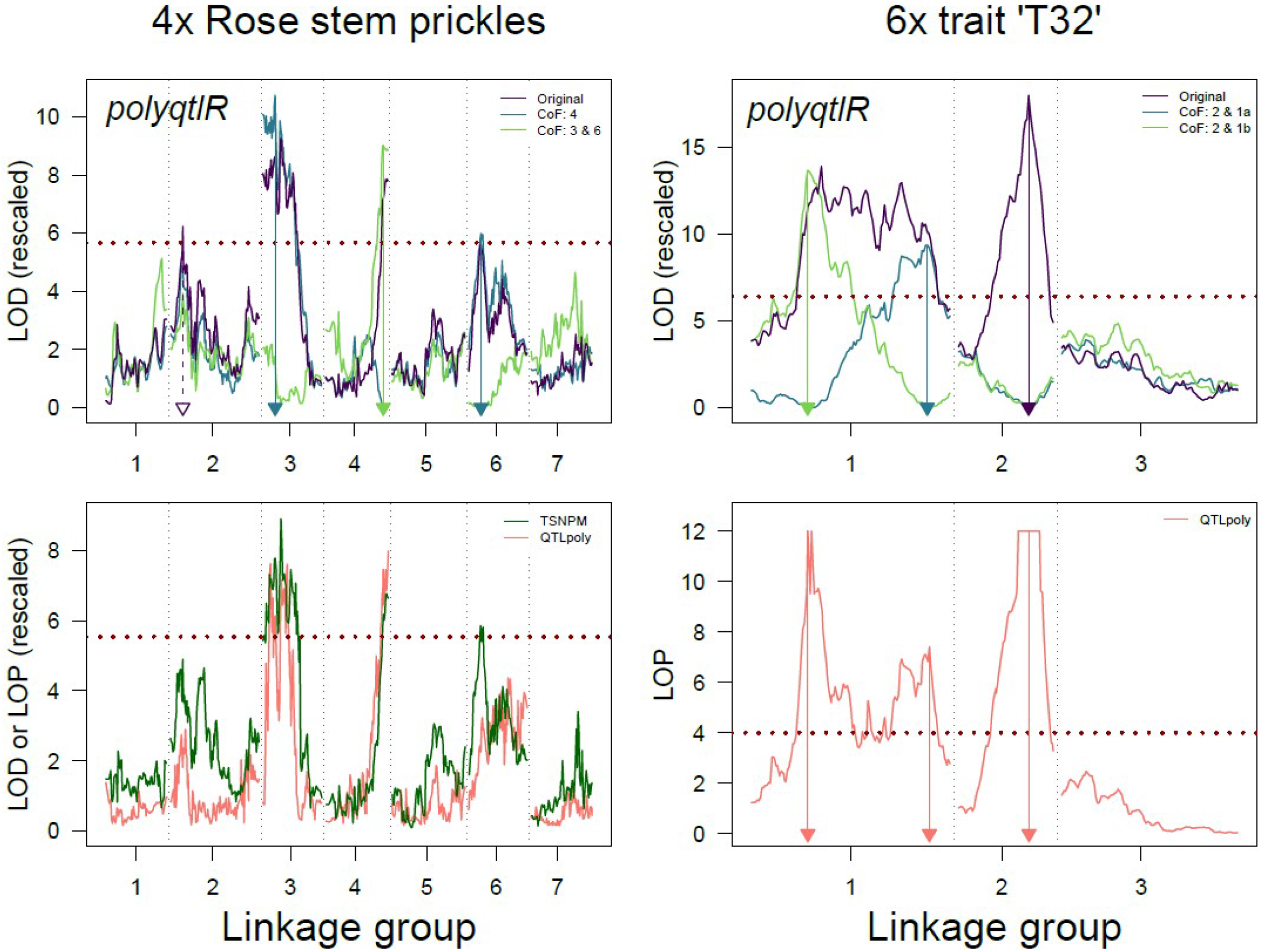
Comparison of the results of *polyqtlR* with those of alternative methods in both a tetraploid and hexaploid dataset. Upper panels: results of *polyqtlR*; Lower panels: results of TSNPM = TetraploidSNPMap and QTLpoly for two example traits: stem prickles in tetraploid rose (left panel) and trait ‘T32’ in a simulated hexaploid dataset (right panel). Estimated QTL positions are highlighted with arrows. In the case of the LG 2 QTL for stem prickles, no significant association was detected after fitting co-factors (dotted purple arrow). Legend “CoF: 3 & 6” refers to a co-factor model with QTL positions on LG 3 and 6 included as co-factors. On the y-axes, LOD or LOP (-log_10_(*p*)) scores were re-scaled so that independently-estimated significance thresholds overlap on the plot. For the trait ‘T32’, QTLpoly returned p-values of 0 around the QTL on LG 2 which cannot be visualised using LOP and were therefore artificially replaced, leading to a plateau around the peak.

For the major QTL detected on LG 3, an additive model with QTL alleles for increased number of prickles located on parental homologues 4 and 6 (*i*.*e*. parental genotypes oooQ x oQoo) was found to have the lowest BIC of the 224 QTL models tested (listed in Supplementary Table1), which corresponded well with the visualised homologue effects for that linkage group (Figure 2).

**Figure 2.**
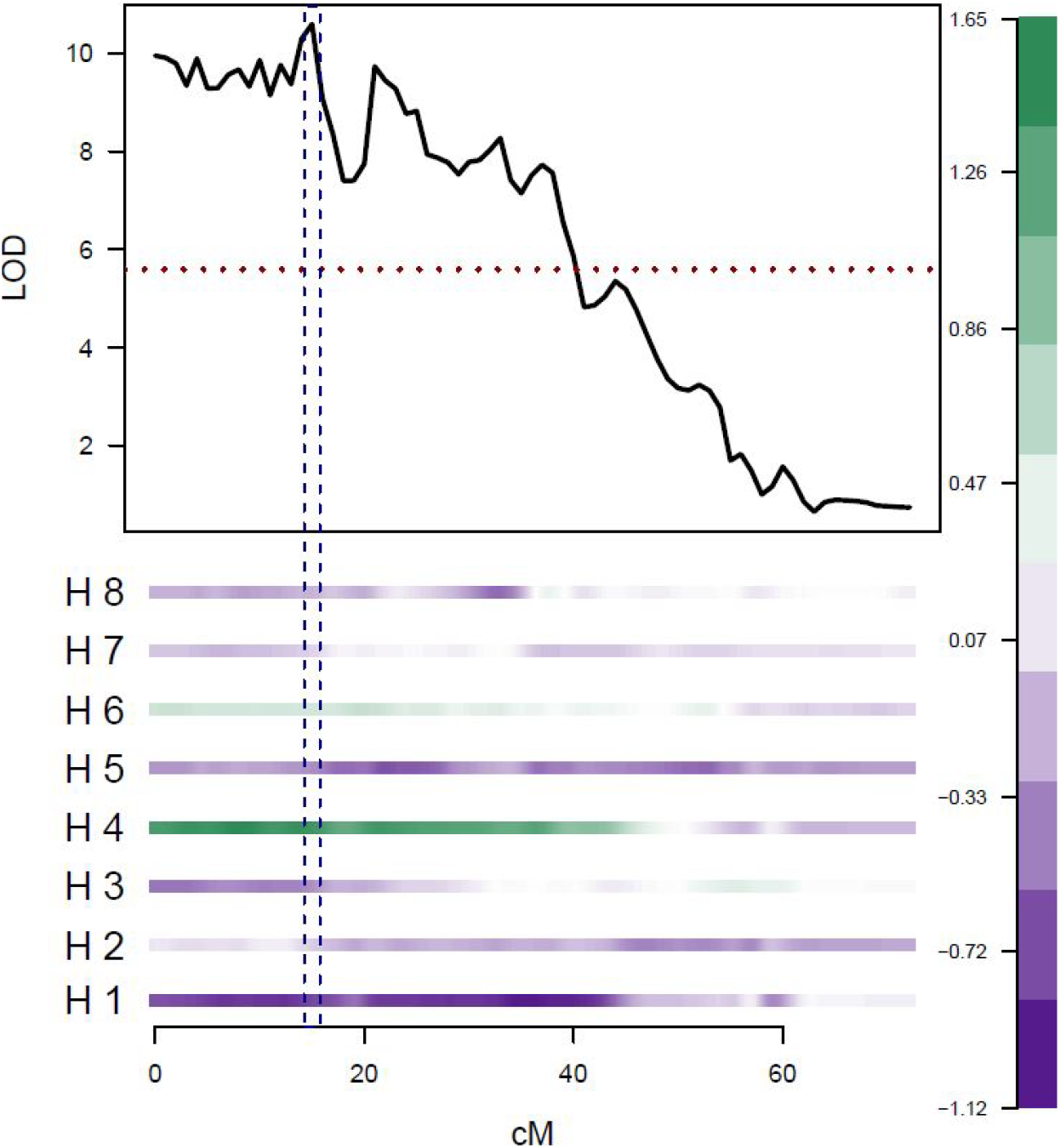
Exploration of the homologue effects at a QTL peak using *polyqtlR*. Example shown corresponds to the QTL peak position on LG 3 of tetraploid rose for the trait stem prickles. Positive effects (increasing the number of prickles) are coloured green, while negative effects are coloured purple. Parental homologues are numbered H1 – H4 (maternal) and H5 – H8 (paternal).

We also used *polyqtlR* and QTLpoly to analyse an example trait ‘T32’ for a hexaploid population (TetraploidSNPMap is restricted to analyses of tetraploid populations and therefore was not included in this comparison). ‘T32’ is a simulated trait provided with the QTLpoly package for test purposes (Supplementary data 1). Two peaks were detected in the initial genome-wide scan using *polyqtlR*, while a third peak became apparent after the major LG 1 peak was fitted as a co-factor (Figure 1). The PVE of the three-QTL model was 50%. The location of the three peaks corresponded very closely to the true positions of the simulated QTL, with all three true QTL positions contained in the LOD – 1 support intervals around the detected peaks. Indeed, *polyqtlR* detected all three simulated QTL with slightly higher precision than QTLpoly: the deviations between peak and true position for QTL 1a, 1b and 2 were 0.97 cM, 7.98 cM and 0.99 cM for *polyqtlR*, while those for QTLpoly were 1.04 cM, 9.26 cM and 1.19 cM, respectively.

### 2. Analysis of meiosis

A hexaploid chrysanthemum (*Chrysanthemum* x *morifolium*) F_1_ population that had previously been used to generate a high-density linkage map (van Geest et al., 2017a) was re-analysed to gain insights into the parental meiosis. An analysis of the genome-wide counts of recombinations across the population showed that the dataset was of remarkably high quality (apart from a pair of outlying individuals), while the parental meioses appear to have involved fewer than the expected average of one cross-over event per bivalent (Figure 3.a).

**Figure 3.**
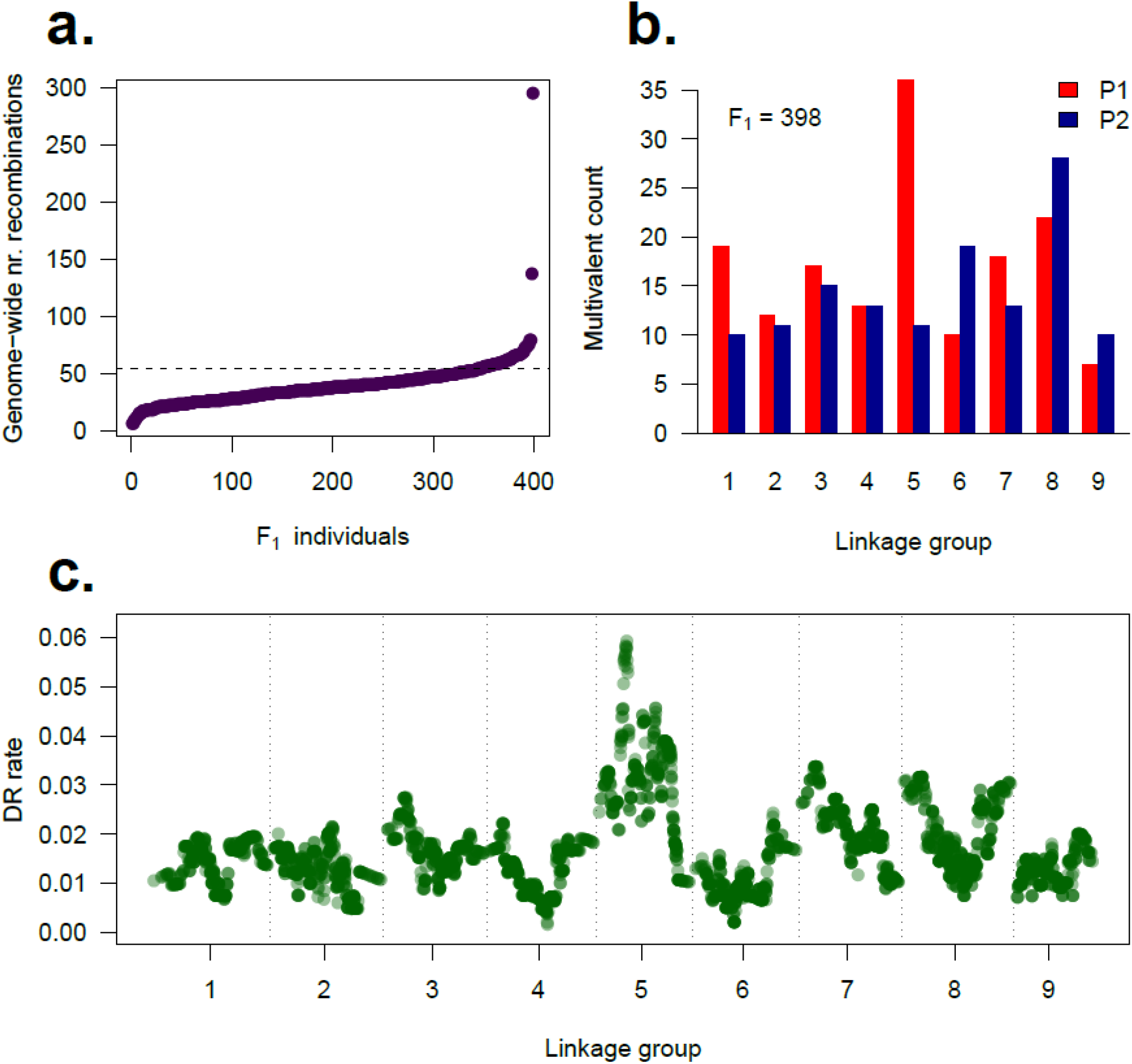
Explorations and visualisations of meiotic dynamics using *polyqtlR*. **a**. Genome-wide counts of predicted recombinations per individual in a hexaploid chrysanthemum F_1_ population, derived from (bivalent-only) IBD probabilities with an error prior of 0.01. The horizontal dotted line shows the expected number of counts assuming on average one cross-over recombination per homologue (6 × 9 = 54); **b**. Numbers of multivalent pairing structures predicted by the HMM per linkage group (x-axis). Maternal counts are shown in red (P1) while paternal counts are shown in blue (P2). Two F_1_ individuals with unusually-high numbers of predicted recombinations in (a) were removed; **c**. Rate of double reduction across the genome, using the same data as (b).

The two outlier individuals were subsequently found to contain significantly more missing values that the rest of the population (Supplementary Figure S1). They were removed and the remaining 398 individuals were re-analysed using the multivalent-aware HMM, allowing the number of multivalents per linkage group to be estimated (Figure 3.b). Maternal LG 5 had an unusually high number of predicted multivalents, which was reflected in a relatively high rate of predicted double reduction events for that chromosome, up to 6 % (Figure 3.c).

With multivalents accounted for, the remaining bivalent pairing configurations were used to test for preferential chromosome pairing. Deviations from a random-pairing (polysomic) model were tested using a chi-square test on the predicted counts of each set of bivalents per homologue (*e*.*g*. a test on the counts of AB, AC, AD, AE and AF for homologue A *etc*.), while the deviations themselves can be used to visualise the chromosomal pairing patterns of both parents using *polyqtlR* (Figure 4). There appeared to be evidence of non-random pairing in the paternal meiosis, with the most extreme deviation identified between paternal homologues H and J of linkage group 1. These homologues were predicted to have paired in 192 of the 389 bivalent-only meioses, an excess of 114 over the number expected if pairing were random (associated chi-square p-values of 2.8 × 10^−44^ and 5.4 × 10^−45^ for homologues H and J, respectively). For a number of chromosomes, three of the fifteen possible bivalent configurations were over-represented, for example in LG 3, 9 (and to a lesser extent LG 4 and 6) of parent 2 (Figure 4). In all such cases (particularly for LG 3 and LG 9), the preferential bivalent pairings were complementary (*i*.*e*. involving all 6 homologues).

**Figure 4.**
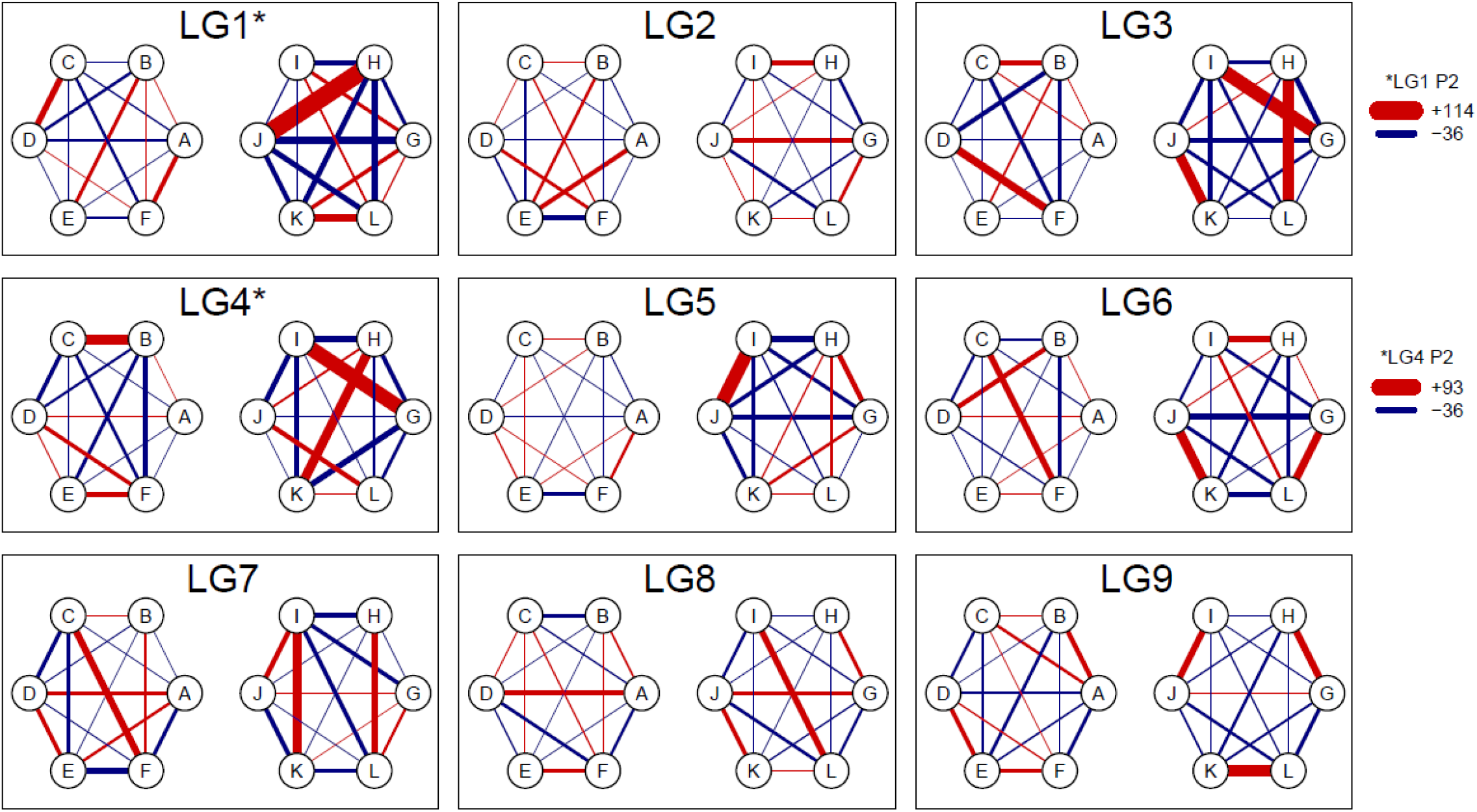
Deviations from a random-pairing model detected in a hexaploid chrysanthemum F_1_ population using *polyqtlR*. Maternal homologues are labelled A - F, while paternal homologues are labelled G – L (these labels are randomly assigned). The thickness of the line connecting parental homologues indicates the level of deviation from a random-pairing model, with counts exceeding expected proportions coloured red, and counts less than the expected proportions coloured blue.

### 3. Speed and accuracy of IBD estimation

The *polyqtlR* package contains two methods to estimate IBD probabilities which are used in many subsequent analyses. We critically compared these methods in terms of their accuracy and computational efficiency. F_1_ populations of 200 offspring each were simulated using PedigreeSim (Voorrips and Maliepaard, 2012) for ploidy levels 2x, 3x, 4x, 6x, 8x and 10x. A range of marker densities were simulated (50, 100, 200, 500, 1000 and 2000 markers per chromosome over 5 chromosomes), as well as differing proportions of simplex x nulliplex markers (proportions from 0 to 1 in steps of 0.2, where “0” contained no 1×0 or 0×1 markers, and “1” contained 50% 1×0 and 50% 0×1 markers). As multivalents were not simulated, these were also not included in the IBD estimation. Computations were performed on a desktop PC (Intel Xeon processor, 3.6 GHz and 16 Gb RAM) in parallel over 5 cores. At lower ploidy levels (2x, 3x and 4x), the HMM was found to be both faster and more accurate than the heuristic method (Figure 5), while at the hexaploid level the HMM was more accurate but had a high computational cost. Hexaploid datasets containing 10,000 markers (2000 per chromosome) were not analysed with the HMM due to protracted run-times. While datasets with higher proportions of simplex x nulliplex markers led to more accurate results using the heuristic method, the opposite was true of the HMM approach (Figure 5). Regardless of the method used, both the error rate and computation time increased with increasing ploidy.

**Figure 5.**
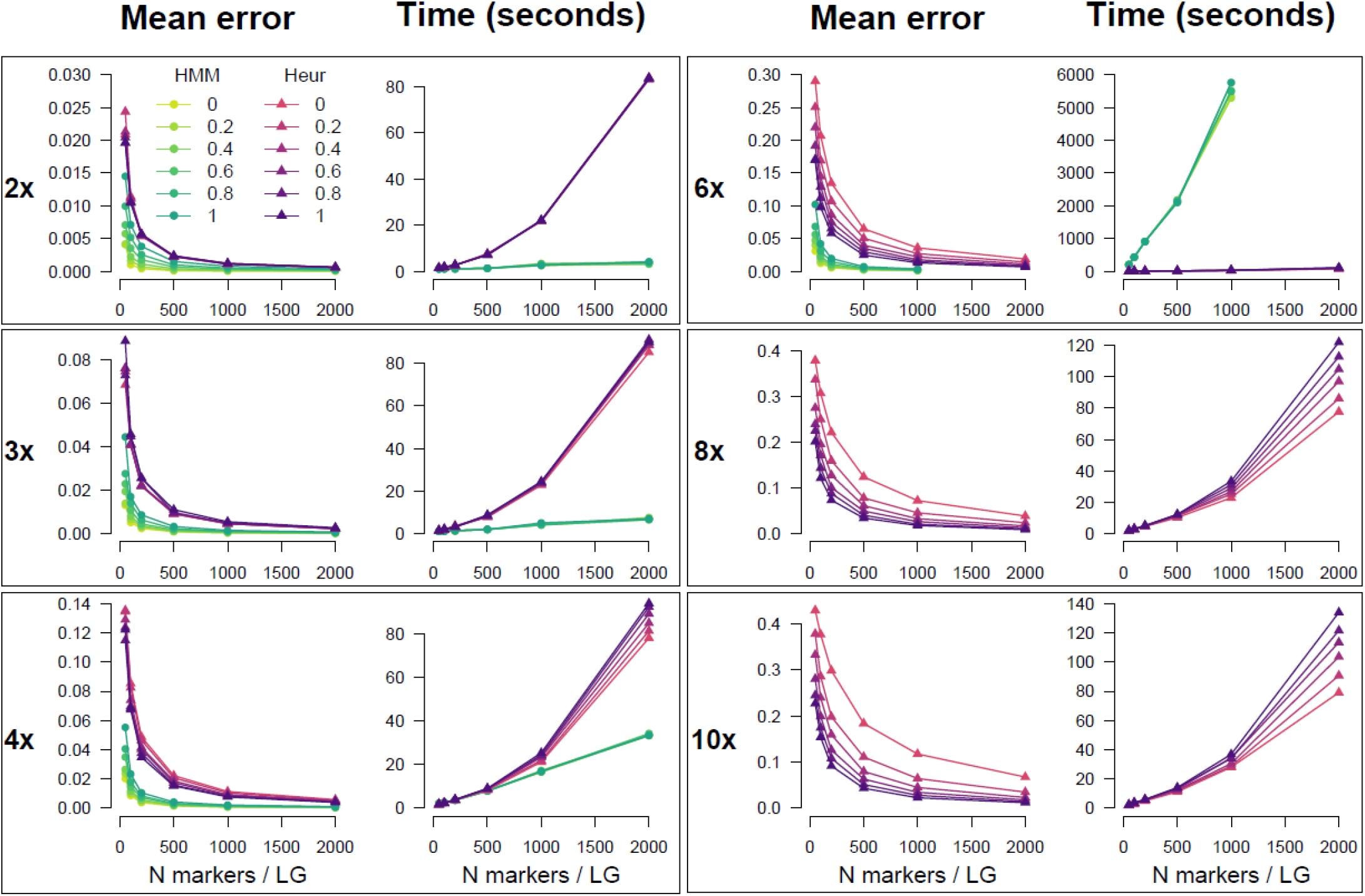
Mean error and computation time associated with estimation of IBD probabilities in *polyqtlR*. Comparison between results from the available options within the package: either a Hidden Markov Model (HMM) or a heuristic algorithm (Heur). In each simulation, 5 chromosomes were simulated for a population of 200 individuals. Mean error was calculated as the average deviation in parental homologue probabilities from the true inheritance probabilities over all estimated positions and individuals. The legend (top left panel) refers to the proportion of simplex x nulliplex markers in the simulated datasets. For higher ploidy levels (8x, 10x), the HMM method has not been implemented and so no comparison was possible.

## Discussion

We are currently witnessing an unprecedented number of developments in polyploid genomics, aided by increasingly affordable genotyping possibilities as well as the realisation among breeders and researchers that genomics-assisted breeding in polyploid species is no longer an insurmountable challenge (Bourke et al., 2018c;Smulders et al., 2019). The *polyqtlR* package aims to facilitate this process by offering a range of tools to help uncover both the origin of favourable (or unfavourable) parental alleles for traits of interest, while also shedding light on how these alleles are passed from one generation to the next by exploring meiotic dynamics of polyploid species.

### Software alternatives

It is usual for new software tools to compare their performance to previously released software alternatives. We have nominally done so by comparing the results of *polyqtlR* to existing packages TetraploidSNPMap and QTLpoly for traits in tetraploid and hexaploid populations, but without quantifying performance differences. The use of the additive-effect interval mapping approach in *polyqtlR* and TetraploidSNPMap is less computationally expensive than fitting mixed models as is done in QTLpoly, and this was indeed reflected in the run-times we observed. For the trait stem prickles in the tetraploid rose population, *polyqtlR* detected four putative QTL, three of which were confirmed by TetraploidSNPMap and two of which were confirmed by QTLpoly and were also detected in independent experiments with diploid rose populations (Crespel et al., 2002;Linde et al., 2006). We feel this demonstrates the importance of comparing results of various software. The peak on LG 2 that we detected may possibly have been a “false positive” detection, although in exploratory analyses these may be less of a concern than possible “false negatives” such as the LG 6 peak that was eliminated in subsequent mapping rounds by QTLpoly. It is interesting to note that the precision of *polyqtlR* for the simulated trait ‘T32’ in the hexaploid population was slightly higher than QTLpoly (all three QTL peak positions were closer to the true QTL positions). This trait has previously been used to demonstrate the superiority of the multi-QTL models over single-QTL ones ((Pereira et al., 2019)), while we have demonstrated here that a “fixed effect interval mapping” approach is equally capable of building an accurate multi-QTL model if genetic co-factors are included. An earlier version of *polyqtlR* was previously used to investigate the effect of double reduction on QTL detection (Bourke et al., 2019). QTL for the traits flesh colour and plant maturity in tetraploid potato coincided with known underlying genes *StCDF1* (Kloosterman et al., 2013) and *StChy2* (Wolters et al., 2010), while a large simulation study confirmed the ability of the package to accurately identify simulated QTL under a wide range of parameter settings (Bourke et al., 2019).

### Hexaploid inheritance

Using *polyqtlR*, we uncovered evidence of preferential pairing in hexaploid chrysanthemum, a phenomenon that was not detected in a previous study using the same population and genotypes (van Geest et al., 2017b)). This highlights the power of leveraging map and genotype information to correctly diagnose preferential pairing, as was done previously in a study of tetraploid rose (Bourke et al., 2017). From our analysis it appears that the hexaploid parents of this population indeed exhibited predominantly hexasomic inheritance, but with some clear exceptions to this trend (Figure 4). Attempting to parametrise preferential pairing at the hexaploid level or higher is clearly non-trivial given the variable patterns of preferential pairing observed here. In some cases, a single pair of homologues behaved as a “sub-genomic unit” (*i*.*e*. showing a strong pairing preference), while elsewhere in the genome, multiple sets of complementary homologue pairs showed non-random pairing patterns, reminiscent of a more allopolyploid-like pairing behaviour for these chromosomes (Figure 4). These sorts of insights could potentially be of enormous importance to breeders aiming to recombine specific alleles on a single homologue (in coupling phase). These meiotic insights are also of fundamental interest to biologists, providing experimental evidence regarding the mode of inheritance of polyploid species that may not fall neatly into the categories of allo- or autopolyploid (Bourke et al., 2017).

### Innovative aspects

*polyqtlR* offers a number of innovations not available elsewhere. For example, it allows the inclusion of multivalent structures in the inheritance model for triploid, tetraploid and hexaploid populations, carried through to subsequent QTL analyses and explorations of parental meioses. At the hexaploid level this is unique, allowing us to estimate rates of multivalent pairing and visualise the double reduction landscape, something that to the best of our knowledge has never previously been visualised in a hexaploid species (Figure 3). The practical implications of double reduction events for QTL mapping may be relatively minor (Bourke et al., 2019), but they can have potentially important breeding implications by increasing the frequency of favourable alleles in particular individuals, as well as being of theoretical interest to polyploid geneticists.

The package also calculates and visualises per-homologue profiles of the genotypic information coefficient (GIC), one of the major factors determining QTL detection power and precision (Bourke et al., 2019). Through visual inspection, parental homologues with poor information can be easily identified and potentially targeted with additional markers.

Options for genotype curation are relatively limited for polyploid species currently, but can be achieved in *polyqtlR* through IBD-informed genotype imputation. The choice of a suitable error prior *ε* in IBD estimation is critical to this step. *ε* is not known *a priori*, but can be estimated *a posteriori* by running the IBD estimation step a number of times with different values (*e*.*g. ε* = 0.01, 0.05, 0.1 and 0.2) and comparing the marginal likelihoods across the mapping population between runs. Applying a higher error prior (*e*.*g. ε* = 0.2) makes transitions between states less probable in the HMM procedure, thus penalising multiple cross-overs that are often necessary to accommodate genotyping errors in a predicted meiotic model with an overly conservative error prior. These can be used directly to re-impute marker genotypes, which could subsequently be used in re-estimating linkage maps that may have been built under the assumption of error-free data.

*polyqtlR* also includes a heuristic approach to IBD probability estimation, something that is not currently available elsewhere but which allows IBD probabilities to be approximated in a relatively short time for populations of *all* ploidy levels, with almost no increase in computation time with increasing ploidy level (Figure 5). Our approach to detecting and visualising preferential chromosome pairing (Figure 4) also provides a clear overview of meiotic pairing and recombination dynamics across experimental populations, leading to insights into pairing behaviour at a level of detail not previously possible. Finally, although not demonstrated here, *polyqtlR* can identify recombinant individuals for specific homologues, a functionality that could be used for tailored breeding approaches or “breeding-by-design” for polyploid crops (Peleman and Van der Voort, 2003).

### Concluding remarks

In this paper we have introduced a novel R package to facilitate QTL analysis and the exploration of chromosomal pairing in polyploid species. *polyqtlR* is freely available under the general public license from the Comprehensive R Archive Network (CRAN) at http://cran.r-project.org/package=polyqtlR.

## Funding

This work was supported by the TKI projects “A genetic analysis pipeline for polyploid crops” (project number TU 263, BO-26.03-002-001), “Novel genetic and genomic tools for polyploid crops” (project number KV 1605-20, BO-26.03-009-004) and the USDA’s National Institute of Food and Agriculture (NIFA) Speciality Crop Research Initiative project “Tools for genomics-assisted breeding in polyploids: Development of a community resource” (2020-51181-32156/SCRI). PMB received an EMBO short term fellowship to work in the group of CAH at BioSS, Dundee, Scotland (ASTF number 228 – 2016). The work of CAH was funded by the Rural & Environment Science & Analytical Services Division of the Scottish Government.

The authors would like to thank Eric van de Weg, Herman van Eck and Giorgio Tumino (WUR Plant Breeding), Michiel Klaassen (Aeres Hogeschool), Camillo Berenos (Dümmen Orange B.V.), Aike Post (Deliflor B.V.) and Jibran Tahir (Plant and Food Research Ltd, New Zealand) for their constructive feedback and discussions.

## Author contributions

Methodology: PMB, REV, CAH, CM; Data analysis: PMB, CAH; Package development: PMB, GVG, JHW; Co-ordination: REV, CM, PA, MJMS, RGFV; Writing manuscript: PMB; Editing manuscript: PMB, REV, CAH, GVG, JHW, PA, MJMS, RGFV, CM. All authors read and approved the final manuscript.

## Supplementary Figures

**Supplementary Figure S1.**
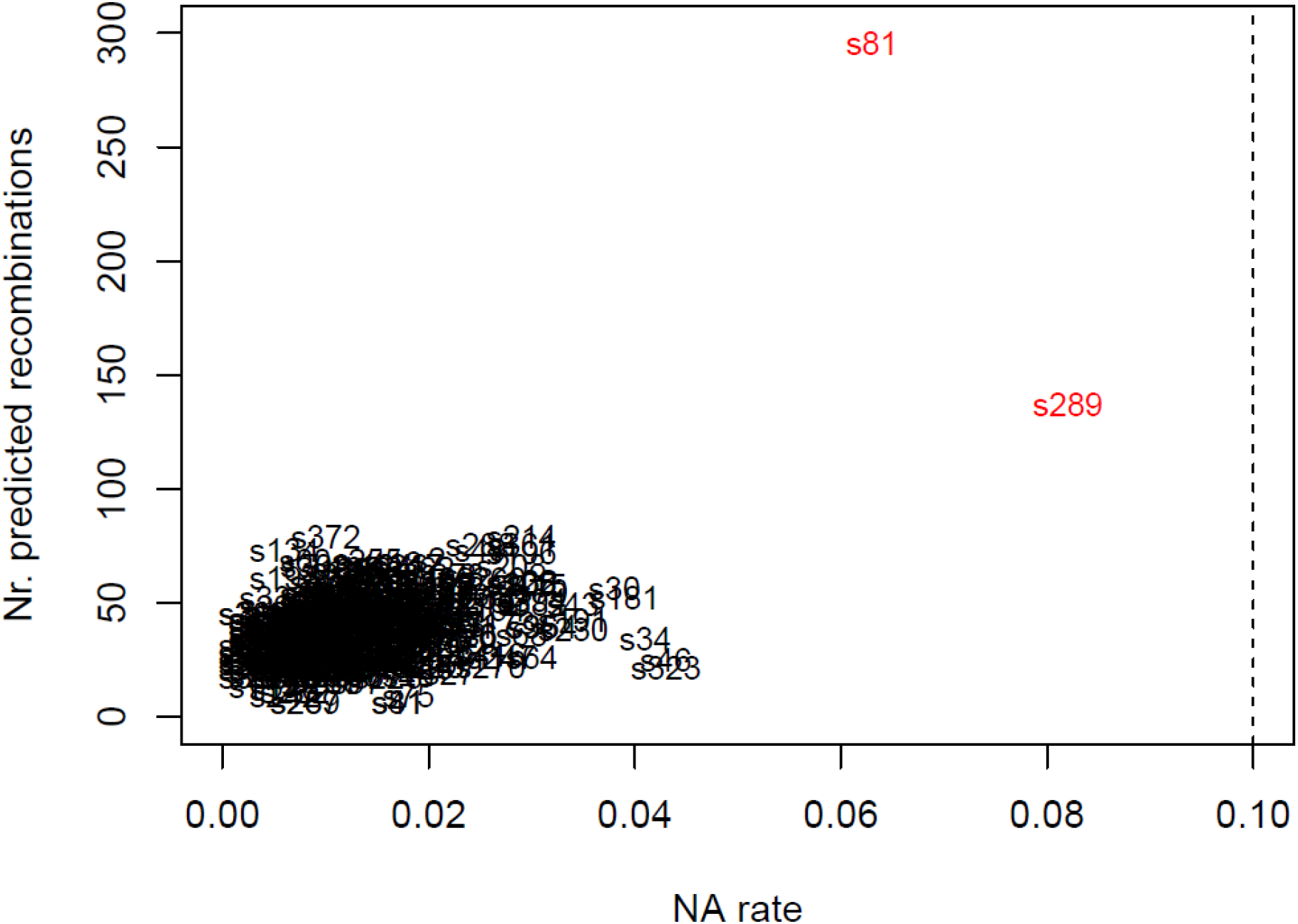
Genome-wide counts of predicted recombinations in a hexaploid chrysanthemum F_1_ population compared with the missing value rates per individual. X-axis shows missing value rates (NA rate). A pair of individuals were identified as having an unusually-high number of recombinations, highlighted in red.

